# Characterization of a Novel Mouse Platelet Transfusion Model

**DOI:** 10.1101/2023.11.10.566577

**Authors:** Dominique Gordy, Theresa Swayne, Gregory J. Berry, Tiffany A. Thomas, Krystalyn E. Hudson, Elizabeth F. Stone

## Abstract

**BACKGROUND:** Platelet transfusions are increasing with advances in medical care. Based on FDA criteria, platelet units are assessed by *in vitro* measures; however, it is not known how platelet processing and storage duration affect function *in vivo*. To address this, we developed a novel platelet transfusion model that meets FDA criteria adapted to mice, and transfused fresh and stored platelets are detected in clots *in vivo*.

**STUDY DESIGN AND METHODS:** Platelet units stored in mouse plasma were prepared using a modified platelet rich plasma collection protocol. Characteristics of fresh and stored units, including pH, cell count, in vitro measures of activity, including activation and aggregation, and post-transfusion recovery (PTR), were determined. Lastly, a tail transection assay was conducted using mice transfused with fresh or stored units, and transfused platelets were identified by confocal imaging.

**RESULTS:** Platelet units had acceptable platelet and white cell counts and were negative for bacterial contamination. Fresh and 1-day stored units had acceptable pH; the platelets were activatable by thrombin and ADP, aggregable with thrombin, had acceptable PTR, and were present *in vivo* in clots of recipients after tail transection. In contrast, 2-day stored units had clinically unacceptable quality.

**DISCUSSION:** We developed mouse platelets for transfusion analogous to human platelet units using a modified platelet rich plasma collection protocol with maximum storage of 1 day for an “old” unit. This provides a powerful tool to test how process modifications and storage conditions affect transfused platelet function *in vivo*.

## INTRODUCTION

Platelets are anuclear discoid cell fragments derived from megakaryocytes, are required for hemostasis, and play crucial roles in inflammation. Platelet transfusions are provided for bleeding prophylaxis in profoundly thrombocytopenic patients and promote clot stabilization in bleeding patients. Approximately 2.5 million platelet units were distributed in 2021, representing a 0.8% increase over 2019 [1, 2]. Notably, 2021 was marked by severe platelet shortages due to the COVID-19 pandemic; advances in chemotherapy, cardiac surgery techniques, and critical care will also likely increase demand for platelet transfusions. There are several FDA-approved platelet preparation and storage methods, along with processing modifications, such as irradiation and pathogen reduction technologies[3]; these may affect platelet function *in vivo*. Given platelet functional plasticity and multiple clinical contexts entailing their use, elucidating how these parameters modulate platelet function is an unmet clinical need.

FDA regulations define platelet unit parameters as: pH ≥ 6.2 at the end of storage, platelet number ≥ 3.0 x 10^11^ (apheresis) or 5.5 x 10^10^ (whole blood), residual white blood cell (WBC) number ≤ 5.0 x 10^6^, and 85% of platelets retained after leukoreduction[4]. Because most platelet units in the United States are stored in plasma at room temperature, they also need to be free of bacterial contamination. Although the gold standard for assessing platelet viability is to measure post-transfusion recovery (PTR) after autologous transfusion of radiolabeled platelet units[5], detecting circulating platelets after transfusion does not measure their functional capacity. Nonetheless, platelet activity in vitro can be assessed by functional assays (e.g., light transmission aggregometry, platelet function analyzer studies, thromboelastography) or by measuring platelet activation using flow cytometry. Using these approaches, numerous in vitro studies suggest that platelet function deteriorates during storage[6-8].

Although many studies examined various aspects of platelet function and transfusion using mouse models, most of these use a modified Tyrode’s solution, with or without other additives (e.g., albumin, apyrase, prostacyclin), to prevent platelet activation during storage[9-14]. However, in the United States, platelet concentrates prepared from whole blood may not be stored in platelet additive solutions, and most apheresis platelet units are also not stored in platelet additive solution[2, 15]. As such, to develop a tractable model to study how platelet processing methods and storage affect platelet function, we developed a reproducible mouse model for transfusing platelets stored in plasma, which satisfies the (adjusted) human FDA platelet transfusion criteria. Our model, which uses a modified pooled platelet rich plasma (PRP) collection technique, provides platelet units with acceptable pH, WBC counts per unit, and PTR. We also developed a novel flow cytometry approach to assess platelet aggregation in vitro, and showed, using confocal microscopy, that these transfused platelets participate in clotting *in vivo*.

## MATERIALS AND METHODS

### Mice

CD-1 (CD-1; strain #022) outbred mice were purchased from Charles River Laboratories. C57BL/6J-TG(UBC-GFP)30Sach/J (green fluorescent protein (GFP) mice; strain #004353) and B6.Cg-Tg(CAG-DsRed*MST)1Nagy/J (red fluorescent protein (RFP) mice; strain #006051) mice were purchased from The Jackson Laboratory. All mice were maintained on a 12:12 light/dark cycle and a standard chow diet (ad libitum). All mouse experiments were approved by Columbia University’s Institutional Animal Care and Use Committee (IACUC).

### Platelet unit preparation

The protocol was modified from the platelet rich plasma (PRP) collection protocols used for human platelets[15]. <u>Fresh frozen plasma (FFP) preparation</u>. To prepare mouse FFP, whole blood (WB) was collected aseptically from GFP mice via cardiac puncture into 15 mL conical tubes with 12.3% CPDA-1 and stored undisturbed at room temperature for 30 minutes. The WB was then centrifuged at 3000 x *g* for 30 minutes at 20°C; all centrifugation steps used the Thermo Scientific Legend XTR centrifuge with an acceleration of 9 and deceleration of 7. The plasma layer was carefully collected, leaving ∼1 mL plasma above the buffy coat (BC) layer so as not to disturb the BC. Plasma was frozen at -20°C in 1 mL aliquots in sterile microcentrifuge tubes. Immediately before use, FFP aliquots were thawed at 37°C and passed through a 0.2 μm filter (Pall, 4612). <u>Platelet unit preparation</u>. On the day of blood collection, WB was collected aseptically from GFP mice via cardiac puncture in 12.3% CPDA-1[16, 17], and the WB sat undisturbed for 30 minutes at room temperature. GFP WB was diluted with sterile PBS at a ratio of 1:2 of WB:PBS, separated into ∼4-6 mL aliquots in 15 mL conical tubes, and then centrifuged at 120 x g for 10 minutes at 20°C. The platelet rich plasma (PRP) and BC with some contaminating red blood cells (RBCs) were placed into new 15 mL conical tubes (hereafter referred to as PRP+BC). PRP+BC was centrifuged at 80 x g for 20 minutes at 20°C, and the resulting PRP was collected carefully without disturbing the BC and RBC pellet. PRP platelets were quantified by flow cytometry (see below), pelleted at 1942 x *g* for 10 minutes at 20°C, and then resuspended in FFP to a final concentration of ∼2.5-3.0 x 10^5^ platelets/μL. Platelet units were stored in mini platelet aliquot bags (BCSI, 6044) with shaking at room temperature.

### Platelet unit analysis and flow cytometry

Platelet units were analyzed for cellular composition (by flow cytometry) and pH, and assessed for bacterial contamination. <u>Flow cytometry</u>. Platelet units were stained with antibodies against CD41a (1:100, eBioMWReg30, PE-Cy7, ThermoFisher), Ter119 (1:1000, APC-efluor 780, ThermoFisher), and CD45 (1:400, 30-F11, Brilliant Violet 605, Fisher) to detect platelets, RBCs, and WBCs, respectively. Data were collected with an Attune NxT flow cytometer with the No-Wash No-Lyse Filter kit (ThermoFisher Scientific; United States) and analyzed with FlowJo software. WBC were quantified using the Leucocount Kit (BD, 340523). Platelet unit pH was measured daily by adding 25 μL of the platelet unit to pH strips with a pH range of 5-9 (Fisher, 13-640-519). <u>Bacterial contamination assessment</u>. Platelet units were cultured at the time of unit preparation, and after 24- and 48-hours of storage, by inoculating pediatric blood culture bottles (BD BACTEC Peds Plus/F, 442020) and incubating them for 1-5 days at 37°C. The culture medium was sampled by adding 1-3 drops to microscope slides followed by Gram stain (BD BBL^TM^ Gram Stain Kits; B12539) and imaging.

### Platelet activation and aggregation

<u>Platelet activation</u> was assessed by incubating 1 μL of the platelet unit with 0.5 U/mL thrombin (Sigma-Aldrich, 604980) diluted in FACS buffer (phosphate-buffered saline + 0.2mg/mL bovine serum albumin + 0.9mg/mL ethylenediaminetetraacetic acid)[18] at room temperature for 2 minutes or with 10mM ADP (Fisher, 22-515-225) at 37 □C for 5 minutes. Platelets were washed with 1 mL FACS buffer, and then stained with t antibodies against CD41a, Ter119, CD45, and CD62P (P-selectin; 1:100, Psel.KO2.3, APC, ThermoFisher). Ter119+ RBCs and CD45+ WBCs were excluded and CD62P expression quantified on CD41a+ platelets. Platelet aggregation was measured by incubating 1 μL of the transfusate (as prepared for platelet PTR; see below) in a total volume of 200 μL with 0.5 U/mL thrombin diluted in FACS buffer at room temperature for 2 minutes or with 10 μL 20mM ADP at 37 □C for 5 minutes. Platelets were washed and stained using the platelet activation protocol. GFP and RFP platelets were evaluated by flow cytometry; platelets doubly expressing GFP and RFP were considered to be aggregated.

### Post-transfusion recovery (PTR)

To determine PTR, a fresh WB unit was prepared from RFP mice. Briefly, WB was collected from 1 RFP mouse by cardiac puncture into CPDA-1 (as above). RFP WB sat for 30 minutes and then was mixed in a 2:1 ratio with a GFP+ platelet unit (fresh, 1- or 2-day stored) to create the transfusate; of note, RBCs in RFP WB do not express RFP (data not shown). Anesthetized CD-1 recipient mice received 150 μL of the transfusate by retro-orbital transfusion. WB was then collected from transfusion recipients by tail prick at 5 minutes, 1, 24, 48, and 72 hours post-transfusion. The ratio of circulating GFP-positive platelets to RFP-positive platelets, relative to this ratio in the transfusate, measured using flow cytometry, was used to determine PTR at each time point.

### Scanning Confocal Microscopy

Transfusates were prepared by mixing GFP platelet units with freshly collected RFP WB in a 3:1 ratio. Anesthetized CD-1 mice were transfused retro-orbitally with 200 μL of fresh, 1-day stored, or 2-day stored transfusate. After 1 hour, a 3 mm section of the tail tip was transected and left to clot. Once clotted, animals were euthanized and 1 cm of the tail tip was transected and sliced longitudinally. Tail tip sections were mounted on microscope slides and imaged by scanning confocal microscopy (Leica TCS SP8, Leica LASX software). The number and percentage of GFP- and RFP-fluorescent platelets were counted and compared to the transfusate composition for each time point. Confocal microscopy images were analyzed by ImageJ software[19].

### Statistical analysis

For comparison of 3 or more groups, a one-way or repeated measures two-way ANOVA with Tukey’s multiple comparisons post-test was utilized; p≤0.05 was considered statistically significant. Statistical analyses were performed using GraphPad Prism.

## RESULTS

### Platelet unit composition and pH change during storage

Per microliter, an average 300mL human platelet unit contains at least 1 x 10^6^ platelets, an average of 1.7 x 10^3^ RBCs[20], and fewer than 16.7 WBCs (<5 x 10^6^ WBCs/unit). Although there is no standard for mouse platelet dose for a platelet transfusion, other groups have published platelet counts ranging from 1.0 x 10^5^ platelets/μL to 4 x 10^6^ platelets/μL[11, 21-24]. Although the final platelet concentration depends on the total volume of FFP into which the PRP is ultimately resuspended, to minimize the mice required for the unit and to allow for sufficient volume for bacterial culturing, in vitro assays, and post-transfusion recovery experiments, we use a minimum platelet count of 1.0 x 10^5^ platelets/μL for the final platelet counts of fresh units.

Mouse platelet units were prepared from GFP mice as described in the Methods section (**Platelet unit Method**, **Fig. 1**). The unit’s platelet count per microliter varied depending on the volume of mouse plasma into which the platelets were ultimately resuspended, but trends between experimental replicates were consistent. For one experimental replicate reported here, fresh platelet units contained the following per microliter: 2.7 x 10^5^ platelets, 680 RBCs, and 6.5 WBCs (**Fig. 2A**). Because platelets have a short half-life, we quantified platelet count as a function of storage age. Intriguingly, platelet counts increased to 3.9 x 10^5^ platelets/μL after 1 day of storage, but significantly decreased below baseline to 8.9 x 10^4^ platelet/μL after 2 days of storage (**Fig. 2B**). Per FDA guidelines, human platelet units require pH ≥ 6.2 [25, 26]. Across all experiments, mouse platelet unit pH ranged from 7.0-7.5 on the day of preparation, 6.4-6.75 on day 1 of storage, and 5.75-6.25 on day 2 of storage (experimental replicate shown in **Fig 2C**.). Platelet unit “swirl,” an indirect measure of platelet discoid shape and suitable pH[27], was observed with fresh and 1-day stored platelets (**supplemental video 1**); platelet swirling was significantly reduced in 2-day stored units. To ensure that platelet units were free from bacterial contamination, samples were incubated in pediatric blood culture bottles. All units tested negative for bacterial contamination during storage (data not shown). These results suggest that this method of preparing mouse platelet units falls in line with modified FDA criteria for human platelets. These mouse platelet units contain sufficient numbers of platelets, are lower than the established thresholds for RBCs and WBCs, and maintain a neutral pH for 1 day of storage. As observed with human platelet units, prolonged storage leads to more acidic pH and reduced viability. Based on these observations, mouse platelet units prepared on the day of collection are considered “fresh” and after 1 day of storage are considered “old.”

**Figure 1.**
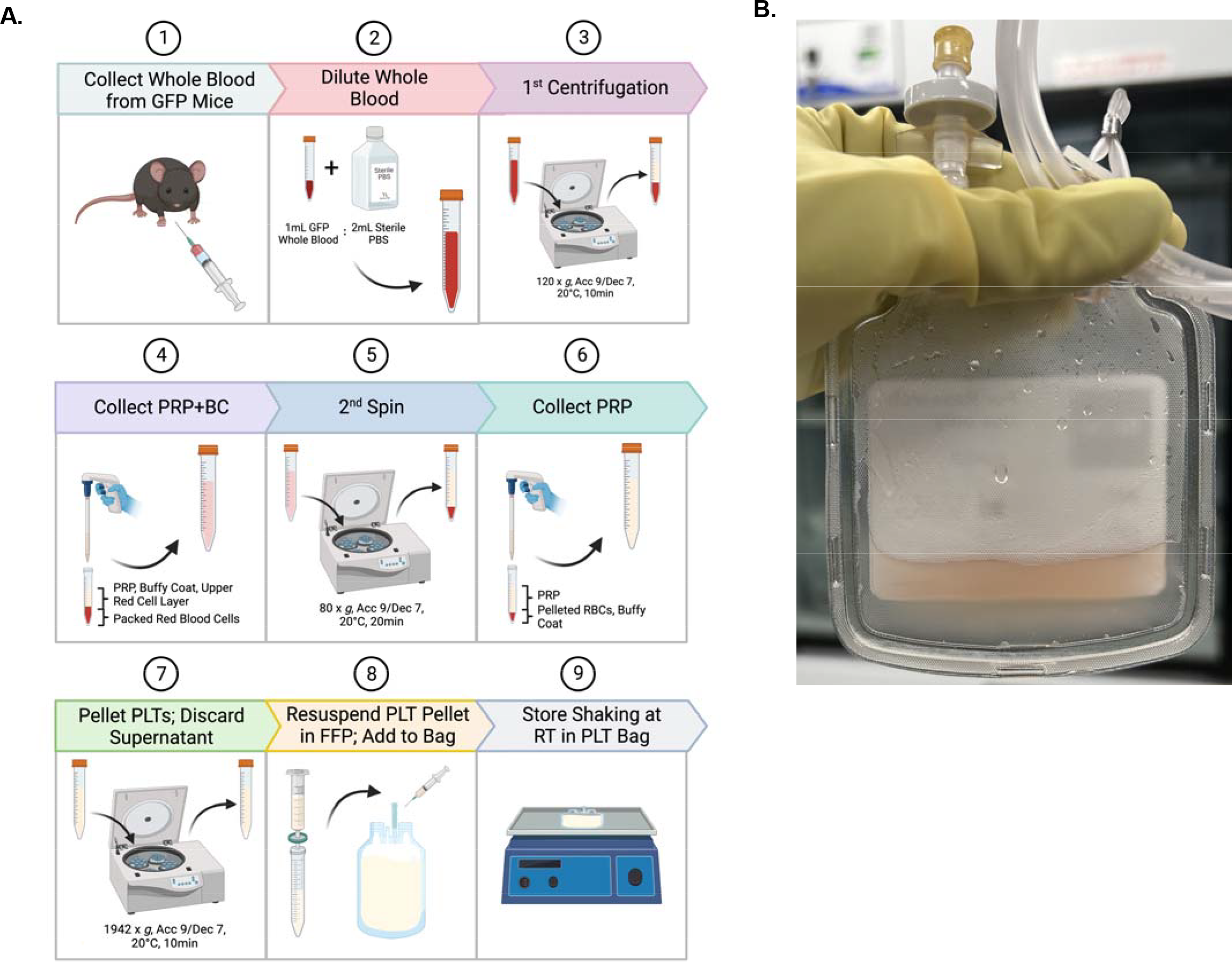
Mouse platelet unit method. (A) Sequential steps to make the mouse platelet unit (Created with BioRender.com). (B) Platelet unit in mini platelet aliquot bag (BCSI, 6044); unit volume is ∼6 mL.

**Figure 2.**
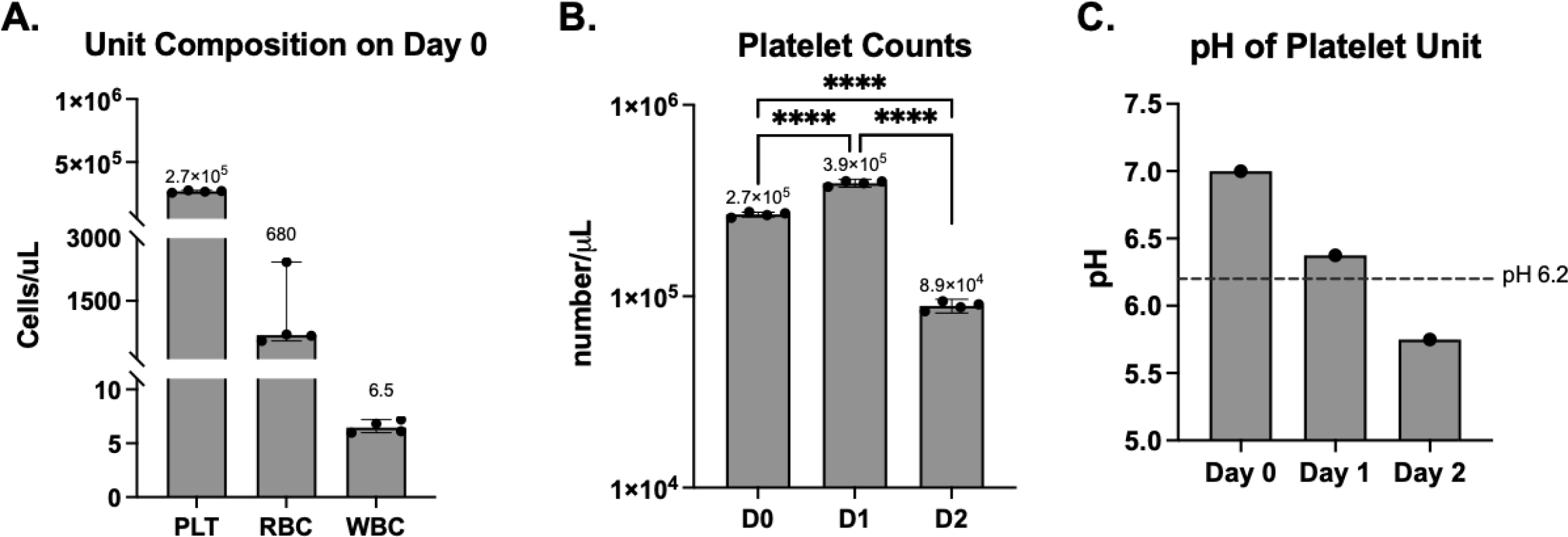
Composition and pH of platelet units change during storage. Aliquots of platelet units were stained with antibodies to detect platelets, RBCs, and WBCs. (A) Freshly manufactured platelet unit contained, on average, 2.7 x 10^5^ platelets (PLT)/μL, 680 RBCs/μL, and 6.5 WBCs/μL. (B) Platelet counts of each unit were calculated throughout storage; platelet count increases to 3.9 x 10^5^/μL after 1 day of storage and decreased to 8.9 x 10^4^/μL after 2 days. (C) Platelet unit pH was tested with pH strips and decreased throughout storage. Dashed line indicates pH 6.2, which is the lower FDA established threshold for human platelet units. Data are representative of 1 of 2 experiments and technical replicates are shown in (A) and (B) with data representing mean +/- SD. Data were analyzed with a one-way ANOVA with Tukey’s multiple comparisons post-test and **** is p<0.0001.

### Storage of platelet units modulates CD62P expression and agonist response

As platelets are often transfused to improve hemostasis and restore clotting capacity, it is essential to test whether platelets become activated upon agonist exposure. To test whether these platelet units could be activated throughout storage, the percentage of platelets expressing CD62P (P-selectin) was quantified at baseline and after incubation with platelet agonists (i.e., thrombin and ADP; **Fig. 3**). Freshly prepared platelet units had low CD62P+ platelet frequencies (mean 13.4%, **Fig. 3B**), which significantly increased throughout storage, demonstrating that platelets were becoming activated during storage. Fresh platelets responded to thrombin and ADP, as measured by significant increases in CD62P+ platelet frequencies (6.3 and 1.4-fold over fresh baseline, respectively). Platelet units stored for 1 day retained functional responses to agonists, as CD62P+ platelet frequencies increased in response to thrombin and ADP (1.85 and 1.6-fold over 1-day baseline, respectively), but responses were less dramatic than seen with freshly prepared platelets. Lastly, 2-day stored platelets had diminished functional responses to thrombin and ADP (1.08 and 1.06-fold over 2-day stored baseline, respectively). Taken together, these data demonstrate that otherwise unmanipulated platelet units become activated throughout the storage duration and lose functional capacity to respond to agonists.

**Figure 3.**
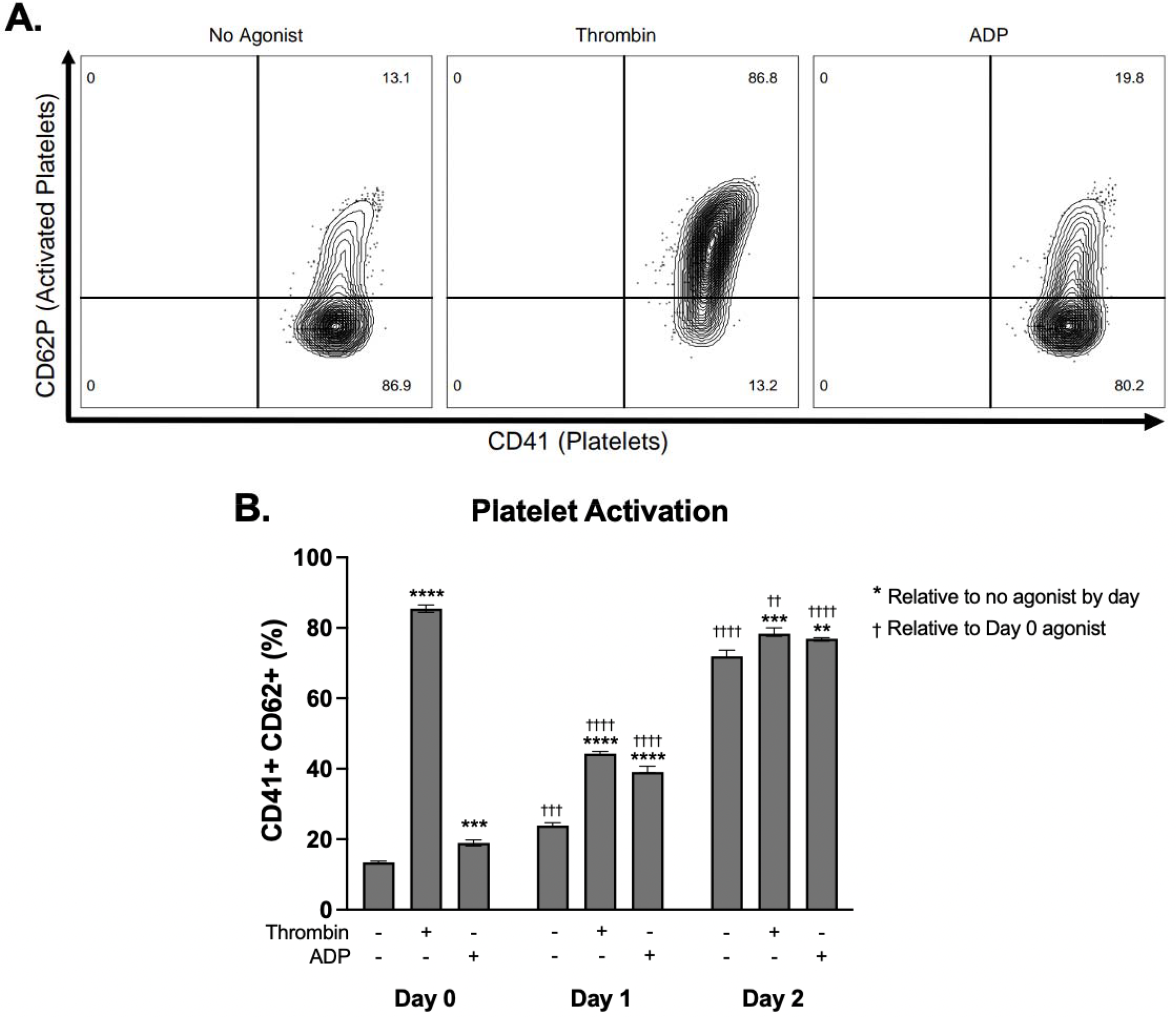
Platelet unit storage modulates CD62P expression and agonist responses. Platelet units were prepared from GFP mice and aliquots from fresh and stored units (1- and 2-day) were stained to detect platelets, RBCs, and WBCs. Ter119+ RBCs and CD45+ WBCs were excluded from the analysis. CD41+ platelets were assessed for CD62P expression. (A) Representative flow plots and (B) frequency of CD41+ platelets in response to no agonist, thrombin (0.5 U/mL), and ADP (10 mM). Data are representative of 1 of 2 experiments and data in (B) are mean +/- SD. Data were analyzed by 2-way ANOVA with Tukey’s multiple comparisons post-test; “*” compares baseline to an agonist by day, whereas “†” compares the agonist condition from day 1 or day 2 to day 0; ** or †† is p<0.01, *** or ††† is p<0.001, **** or †††† is p<0.0001.

### Platelet aggregation diminishes with prolonged storage

Another measure of platelet function is the ability to aggregate and form clots. As conventional aggregometry assays require large volumes of blood (i.e., 800 μL), we adapted a recently-reported assay [28], to develop a new method using flow cytometry that requires smaller volumes. To that end, we avoided ex vivo platelet labeling by using platelets collected from two strains of transgenic mice with platelets expressing either GFP or RFP, thereby enabling visualization of platelet aggregation in vitro by flow cytometry. As such, individual platelets express GFP or RFP and platelet aggregates are positive for both GFP and RFP (**Fig 4A**, representative flow plots).

**Figure 4.**
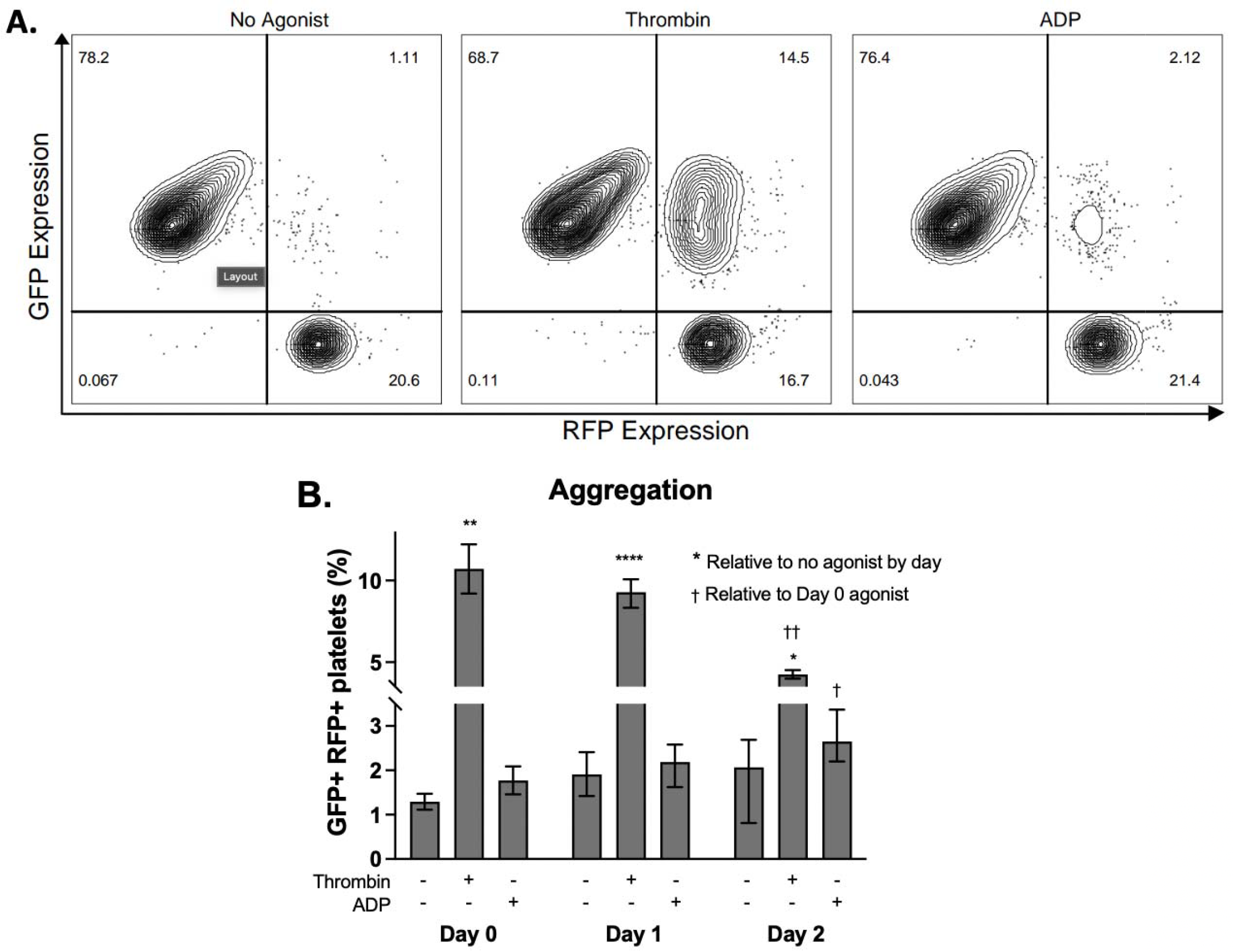
Platelet aggregation diminishes with prolonged storage. GFP platelet units after different times in storage were mixed with fresh RFP whole blood (WB), incubated in FACS buffer with thrombin (0.5 U/mL) or ADP (10 mM), and then analyzed by flow cytometry for the percentage of GFP+RFP+ (i.e., double positive) platelet “clumps,” signifying platelet aggregation. (A) Representative flow plots of fresh GFP platelet units and fresh RFP WB exposed to vehicle, thrombin, and ADP. (B) Percentage of GFP+RFP+ platelet aggregates formed from GFP units (fresh, 1-day or 2-day stored) mixed with fresh RFP WB and unstimulated or activated with thrombin or ADP. Data shown are representative of 1 of 2 experiments and data represent mean +/- SD. Data were analyzed with by 2-way ANOVA with Tukey’s multiple comparisons post-test and “*” compares baseline to an agonist by day, whereas “†” compares the agonist condition from day 0 to day 1 or day 2 (* or † is p<0.05; ** or †† is p<0.01; **** is p<0.0001).

To test how our platelet preparation affected aggregation, GFP platelet units (fresh or stored) were mixed with fresh WB from RFP animals (which have RFP platelets) followed by agonist exposure. Fresh GFP platelets mixed with RFP WB had few detectable aggregates (**Fig. 4A**, **left and 4B**), suggesting that these platelets are not activated. To induce activation and promote aggregation, fresh platelets were treated with thrombin (**Fig. 4A**, **middle**) or ADP (**Fig. 4A**, **right**); thrombin-treated, but not ADP-treated, fresh platelets displayed significant increases in GFP+RFP+ frequencies, as compared to the “no agonist” control (**Fig. 4B**). Similar to fresh GFP units, thrombin treatment, but not ADP, led to significant increases in aggregation of 1-day stored GFP units, as compared to the “no agonist” control. A similar trend was observed with 2-day stored GFP platelet units, although aggregation frequencies were diminished. Taken together, these data show low levels of platelet aggregation in otherwise unmanipulated platelet units throughout a 2-day storage period. Additionally, thrombin exposure consistently induced platelet aggregation, but its effect diminished throughout storage. Lastly, platelet exposure to ADP did not promote significant aggregation. In sum, these data, along with those in **Fig.3**, suggest that this platelet processing procedure and storage conditions yield a functional platelet product by in vitro metrics.

### PTR of fresh and stored platelet units

To test whether platelets from these units circulated *in vivo*, GFP platelets (fresh or stored) were mixed with RFP WB at a ratio of 2:1 and transfused into wild-type mice (**Fig. 5A**). Peripheral blood was collected at defined time points and the GFP:RFP ratio was normalized to the percentage of GFP+ platelets in the transfusate to determine PTR. Fresh and 1-day stored platelets had similar PTRs over the first 24 hours post-transfusion; diversion of PTRs was evident at 72 hours post-transfusion but both remained above 60% (**Fig. 5B**). In contrast, 2-day stored platelets had a PTR under 60% at the first time point (i.e., 5 minutes). Taken together, these data demonstrate that fresh and 1-day stored platelet units had excellent PTR, even at 72 hours post-transfusion, but 2-day stored platelets do not circulate well post-transfusion.

**Figure 5.**
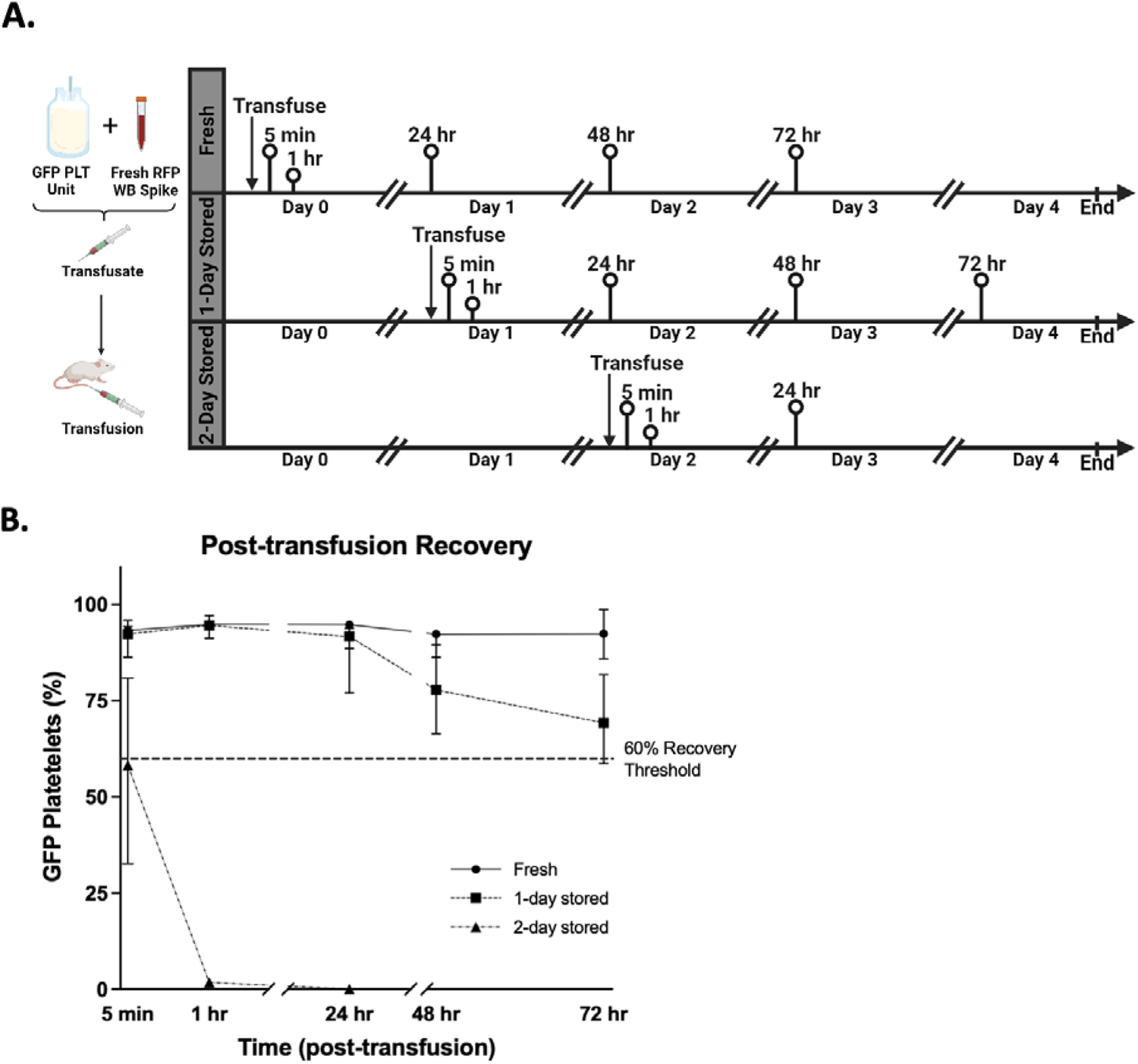
Post-transfusion recovery (PTR) of fresh and stored platelet units. (A) Schematic of PTR protocol. GFP platelet units stored for different durations (fresh, 1-day stored, 2-day stored) were mixed 2:1 with fresh RFP WB and 150μL were transfused retro-orbitally into recipient CD-1 mice (n = 5 per group). Post-transfusion recovery was quantified at different times post-transfusion by flow cytometry. Created with BioRender.com. (B) Graph of PTR for fresh, 1-day stored, and 2-day stored GFP platelet units. The percentages of GFP platelets were normalized to the composition of the transfusate from each day transfused. Fresh and 1-day stored platelet units had a PTR of >60% even 72 hours after transfusion, whereas 2-day stored units had very poor PTR, with only 2% GFP platelets circulating at 1 hour, and no GFP platelets circulating by 24 hours.

### Transfused GFP platelets from fresh and 1-day stored units participate in clotting

To determine whether transfused platelets contribute to clot formation *in vivo*, recipient mice were transfused with a 3:1 ratio of GFP platelet units (fresh or stored) and fresh RFP WB, which served as a loading control. Post-transfusion, a 3mm section of the tail tip was transected, and tails permitted to clot. Following euthanasia, tail tips were imaged by scanning confocal microscopy (**Fig. 6A**). The number of GFP and RFP platelets in each clot from each recipient were counted and the frequency of GFP platelets in the clot calculated; this frequency was compared to the GFP frequency in the transfusate (**Fig. 6B**). Of note, significantly fewer GFP platelets from 2-day stored units were detected in clots, as compared to fresh and 1-day stored platelets. However, to assess whether a disproportionate frequency of GFP platelets was actually detected in clots, as compared to the transfusate, GFP frequencies were normalized (**Fig. 6C**); GFP platelets from fresh and 1-day stored units were detected in clots at similar frequencies, but 2-day stored platelets were significantly reduced. Taken together, these data show that transfused fresh and 1-day stored GFP platelet units participate in clotting in this tail transection model, whereas transfused 2-day stored GFP platelets are not found in these clots.

**Figure 6.**
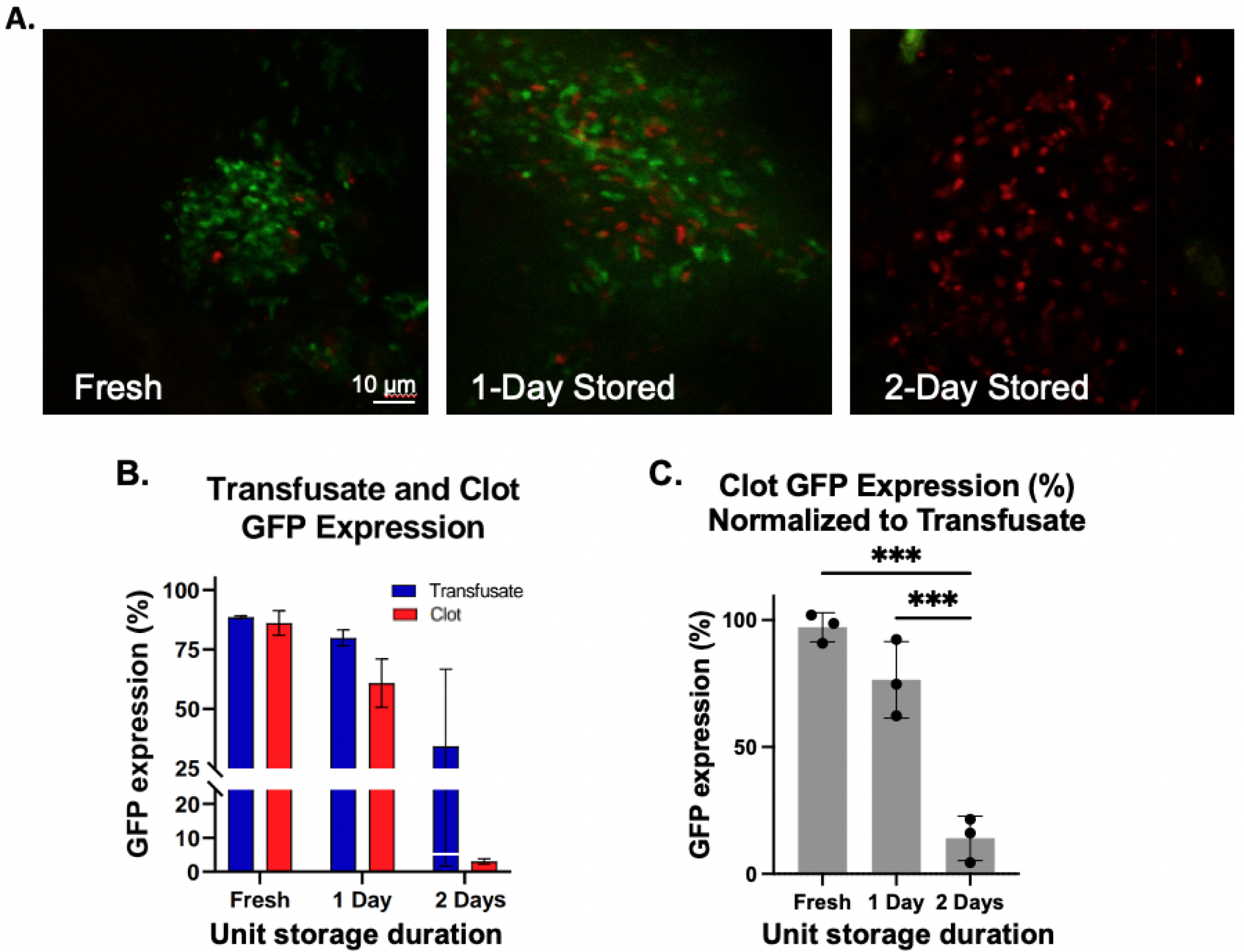
Transfused GFP platelets are identified in clots using fresh and 1-day stored platelet units but not 2-day stored platelet units. Recipient mice were transfused with a 3:1 transfusate of fresh or stored GFP units mixed with fresh RFP whole blood, followed by tail transection and mounting of tail tips for multiphoton microscopy. (A) GFP and RFP platelet contributions to the clots from fresh, 1-day stored, and 2-day stored platelet units. Bar, 10 μm. (B) Graph comparing levels of GFP-expressing platelets as a percentage of total fluorescence (GFP + RFP) in the transfusate (blue bars) and clot (red bars); the percentage of GFP in the transfusate compared to clots for each day of storage was not significant. (C) When normalized to the percentage of GFP-expressing platelets in the transfusate by day of storage, the percent of GFP expression in the clot was significantly reduced for platelet units stored for 2 days (*** p<0.001). Data shown are cumulative from 3 independent experiments, with n=2 recipients per group. Data were analyzed with a one-way ANOVA with Tukey’s multiple comparisons post-test and *** is p<0.001.

## DISCUSSION

We present a model for mouse platelet transfusion involving storing mouse platelet concentrates in plasma with sufficient platelet counts and acceptably low numbers of RBCs and WBCs. Fresh and 1-day stored platelets in these units activate in vitro, aggregate in vitro and *in vivo*, exhibit excellent PTR at 72 hours, and participate in clotting. Additionally, this mouse model is highly reproducible between experimental replicates.

There are several similarities and differences between this mouse model and human platelet units. In our hands, we found that diluting WB in PBS before centrifugation allows for better separation of PRP, which was previously published [14, 22]. We then pellet and resuspend the mouse platelet units in FFP, instead of fresh donor plasma, to reduce the time required for platelet unit manufacturing. Surprisingly, performing the serial centrifugation steps without leukoreduction resulted in fewer WBCs per microliter in the final product, as compared to leukoreduction using a neonatal leukoreduction filter (data not shown). Although this deviates from the protocol for manufacturing human platelet rich plasma, this centrifugation without leukoreduction led to less manipulation of the unit. Additionally, the platelet counts in the mouse units increased between fresh and 1 day of storage; this is also seen with human platelet units manufactured from whole blood, presumably because small platelet clumps disaggregate during storage with agitation[29]. Lastly, in the mouse model, platelet units that have been stored for 2 days have low pH, and the resulting platelets do not circulate or participate in clot formation. Thus, in this model, when mouse platelet units are stored at room temperature with agitation, the limit of storage is 1 day.

In summary, this mouse model for manufacturing platelet concentrates stored in plasma takes advantage of GFP and RFP transgenic lines. Using this model, the effects of various storage modifications, including irradiation, pathogen reduction technologies, platelet additive solutions, and cold storage, can be studied using relevant *in vivo* and in vitro outcome measures. This model will also provide a powerful tool to interrogate important clinical scenarios, including platelet function during intracerebral hemorrhage and platelet alloimmunization, as this murine model effectively recapitulates human transfusion.

## ATTRIBUTION

TAT, KEH, and EFS designed the studies and the experiments. DG collected and analyzed data from mouse experiments. DG, TS, and ES performed confocal microscopy, and DG and GJB performed microbiology analysis of the platelet units. All authors were involved in the interpretation of data. KEH and EFS wrote the manuscript. All authors contributed to the manuscript and approved the submitted version.

## ACKNOWLEDGEMENTS

The authors acknowledge that these studies used the Confocal and Specialized Microscopy Shared Resource of the Herbert Irving Comprehensive Cancer Center at Columbia University, funded in part through the NIH/NCI Cancer Center Support Grant P30CA013696.

